# Ms4a4a loss preferentially remodels microglia with limited, compartment-specific effects across central and peripheral immune compartments in 5xFAD mice

**DOI:** 10.64898/2026.10.26.762005

**Authors:** Emma P. Danhash, Anthony C. Verbeck, Abhirami K. Iyer, Celeste M. Karch

## Abstract

**Background:** Genome-wide association studies have identified the *MS4A* locus as a major modifier of Alzheimer’s disease (AD) risk and resilience. *MS4A4A* is enriched in microglia, and we previously demonstrated that loss of *Ms4a4a* reduces amyloid plaque pathology and attenuates amyloid-associated tau spreading. However, *MS4A4A* is also expressed in peripheral myeloid populations, raising the possibility that its effects extend beyond brain-resident microglia. Here, we sought to define how constitutive *Ms4a4a* loss alters immune populations across central nervous system (CNS), CNS-border, and peripheral compartments in the context of amyloid pathology.

**Methods:** We performed multicompartment immune profiling by flow cytometry in 5xFAD *Ms4a4a* wild-type and 5xFAD *Ms4a4a*-deficient mice. Myeloid and lymphoid populations were characterized in the brain, dural meninges, skull bone marrow, tibial bone marrow, and spleen.

**Results:** *Ms4a4a* loss produced its strongest immune effects in brain microglia. 5xFAD *Ms4a4a*-deficient mice exhibited increased microglial abundance, increased frequency and expression of CD86, increased MHC class II expression intensity, and enrichment of CXCR4-high microglia, whereas P2RY12 and TREM2 were unchanged. Brain macrophages showed increased MHC class II expression, whereas conventional dendritic cells (cDCs) in the brain showed reduced MHC class II expression, with limited changes in other infiltrating immune populations. Immune composition in the dura and skull bone marrow was largely preserved, although MHC class II expression was increased in skull bone marrow cDCs. Peripheral immune composition in tibial bone marrow and spleen was similarly preserved, with selective increases in M2-like macrophages in tibial bone marrow and splenic CD8 T-cell frequency.

**Conclusions:** These findings identify microglia as the immune population most strongly remodeled by *Ms4a4a* loss in the setting of amyloid pathology. Constitutive *Ms4a4a* deficiency does not produce broad systemic immune remodeling but instead preferentially alters microglial state, accompanied by more limited, cell type- and tissue-specific effects across other central and peripheral immune compartments. These findings establish MS4A4A as a context-dependent regulator of immune responses to AD pathology with particularly prominent effects on microglia.

## Introduction

Genome wide association studies have identified numerous loci associated with Alzheimer’s disease (AD) risk, with many risk genes enriched in microglia and other myeloid cells [1–12]. These findings have established immune pathways as important contributors to AD pathogenesis. While microglia are the primary resident immune cells of the brain, immune responses relevant to AD are not restricted to the brain parenchyma. Peripheral myeloid and lymphoid cells can enter the AD brain and influence local inflammatory responses and Aβ clearance, with the potential to promote or limit pathology depending on disease stage and context [13–23]. The dural meninges and skull bone marrow form the CNS-border immune compartments that communicate with the brain, shape neuroinflammatory responses, and are remodeled in AD [24–29]. Alterations in immune populations have also been reported in the spleen and peripheral bone marrow in mouse models of AD [26, 30–35]. Thus, defining whether AD-associated immune genes act broadly across these compartments or preferentially within specific immune populations is important for understanding their contribution to disease.

The *MS4A* locus has emerged as a major modifier of AD risk and resilience [8–10, 36–38]. Within this locus, *MS4A4A* is highly enriched in microglia, and genetic variation near *MS4A4A* is associated with AD risk and cerebrospinal fluid levels of soluble TREM2 (sTREM2) [6, 36, 37, 39]. However, *MS4A4A* expression is not restricted to microglia. In peripheral tissues, *MS4A4A* is expressed in monocytes, macrophages, and dendritic cells [40, 41]. Functional studies have implicated MS4A4A in TREM2- and Dectin-1-associated signaling, macrophage activation states, and tissue-resident macrophage responses [42–48]. Together, these observations raise the possibility that MS4A4A regulates immune responses in a cell type- and tissue-dependent manner.

We previously demonstrated that constitutive loss of *Ms4a4a* modifies AD-related pathology in 5xFAD mice. *Ms4a4a* deficiency increases amyloid beta (Aβ) clearance, reduces amyloid plaque burden, and promotes plaque compaction, while altering microglial inflammatory state [49]. We further found that *Ms4a4a* loss attenuated dense core plaque-associated tau spread [50]. Consistent with these findings, recent studies have implicated *MS4A4A* in microglial transcriptional programs associated with inflammatory signaling, autophagic and lysosomal function, and proteostasis [47, 50]. Together, these studies support a role for MS4A4A in regulating microglial responses to AD pathology. However, because these studies used constitutive *Ms4a4a* loss, it remains unclear whether the resulting immune consequences of *Ms4a4a* deficiency are predominantly microglial or reflect broader remodeling across other central and peripheral myeloid and lymphoid immune cell populations.

Here, we performed multicompartment immune profiling by multiparametric flow cytometry to define the impact of constitutive *Ms4a4a* loss on immune populations in the brain, dural meninges, skull bone marrow, tibial bone marrow, and spleen in the context of amyloid accumulation. *Ms4a4a* deficiency produced its strongest effects in brain microglia, increasing microglial abundance and selectively altering CD86, MHC class II, and CXCR4-associated pathways. Other brain, CNS-border, and peripheral immune populations exhibited more limited and cell type-specific changes. Together, these findings identify microglia as the prominent immune population remodeled by *Ms4a4a* loss while demonstrating that its effects across the broader immune system are selective rather than uniform.

## Methods

### Animals

Animal care and surgical procedures were approved by the Animal Studies Committee of Washington University School of Medicine (Protocol #23-0237) in accordance with guidelines from the United States National Institutes of Health.

5xFAD (B6.Cg-Tg (APPSwFlLon, PSEN1*M146L*L286V) 6799Vas/Mmjax, Stock number 034848-JAX, MMRRC, The Jackson Laboratory) hemizygous mice were crossed with *Ms4a4a^+/-^*mice (generously provided by Alector; constitutive knockout generated using CRISPR/Cas9 to excise exons 1-7 in C57BL/6 mice) to generate 5xFAD x *Ms4a4a^+/-^*mice [49]. Next, 5xFAD x *Ms4a4a^+/-^*were crossed with Ms4a4a^+/-^mice to obtain 5xFAD x *Ms4a4a^-/-^*(5xFAD 4A-KO; n = 5) mice and 5xFAD x *Ms4a4a^+/+^* (5xFAD 4A-WT; n = 8) littermate controls. Genotype was confirmed using primers to determine the presence of *Ms4a4a* and the 5xFAD transgene (**Supplemental Table 1**). Six-month-old female mice were used in this study.

### Retro-orbital injection of APC/Cy7-CD45 antibody and tissue collection

At 6 months of age, 5xFAD 4A-KO and 5xFAD 4A-WT mice were anesthetized with isoflurane and retro-orbitally injected with an APC/Cy7-CD45 antibody (100uL; 3mg/mL). Five minutes after injection, mice were anesthetized with sodium pentobarbital and spleens were removed and placed in RPMI 1640 (Gibco, 11875093) supplemented with 10% FBS, L-glutamine, and penicillin-streptomycin (RPMI+). Mice were then perfused with ice cold PBS.

Skulls were cut along the parietal ridge from the foramen magnum to the olfactory bulbs [51]. Skullcaps with attached dural meninges were temporarily stored in RPMI+ on ice. Brains were harvested, and the cerebellum and olfactory bulbs were removed. Brains were temporarily stored in Dounce Buffer (1X HBSS, 5 mM HEPES, 0.5% glucose) on ice. A single tibia was extracted from each mouse and stored in RPMI+ on ice. All attached tissue was cleaned off bones.

### Tissue preparation for flow cytometry

Brains were processed for flow cytometry using a previously described method with all steps completed on ice [52]. Briefly, using a razor blade, brains were chopped to roughly 1mm^3^ pieces then transferred to a 5mL Dounce homogenizer containing 5mL of Dounce Buffer. Tissue was homogenized with 6 to 10 full strokes until the solution appeared to be homogenous then passed through a 40µm strainer into 50mL tubes. Samples were centrifuged at 500 ×g for 5 minutes at 4°C with the brake set to 5 and resuspended in 30% Percoll in RPMI 1640 with 2mL of 1X Dulbecco’s phosphate-buffered saline carefully overlaid. Samples were centrifuged at 400 ×g for 20 minutes at 4°C with the brake set to 0. The myelin layer and supernatant were removed, and the remaining cell pellet was resuspended in RPMI 1640. A cell count was then taken using a Countess 3 (Invitrogen), and the cell suspension was stored on ice until staining for flow cytometry.

Dural meninges were peeled off skullcaps and processed as previously described with modifications [51]. Briefly, dural meninges were enzymatically dissociated in pre-warmed dissociation buffer containing RPMI 1640, collagenase VIII (1mg/mL, Millipore Sigma, DN25-100MG), DNase I (0.5mg/mL, Millipore Sigma, C2139-100MG), and FBS (2% v/v, Biowest, S1620) for 20 minutes, with five triturations using a P1000 micropipette at after 10 minutes of incubation. RPMI supplemented with FBS (10% v/v) was added to neutralize the enzymatic reaction, and cell suspensions were filtered through a 70µm strainer. Samples were centrifuged at 450×g for 5 minutes at 4°C, resuspended in FACS buffer and stored on ice until staining.

Bone marrow suspensions were prepared as previously described [53]. Skull bone marrow was isolated by mechanically dissociating the skull in PBS using scissors followed by crushing with a pestle. Tibial bone marrow was flushed out of the bone with RPMI+. Skull and tibial bone marrow were filtered through a 70µm strainer and centrifuged at 450×g for 5 minutes at 4°C. Samples were then resuspended in Red Blood Cell Lysing Buffer (Millipore Sigma, R7757-100ML) and incubated for 2 minutes at room temperature. RPMI+ was added to quench the reaction, after which cells were centrifuged at 450×g for 5 minutes at 4°C, resuspended in FACS buffer, and stored on ice until staining.

Spleens were processed using a previously described protocol with minor modifications [54]. Briefly, spleens were gently pressed through a 40µm strainer positioned over a 50mL conical tube using a clean syringe plunger, after which the strainers were washed with RPMI 1640. Samples were centrifuged at 600×g for 7 minutes at 4°C. Cells were resuspended in Red Blood Cell Lysing Buffer and incubated for 2 minutes at room temperature. RPMI+ was added to quench the reaction, and samples were then centrifuged at 600×g for 7 minutes at 4°C. Pellets were resuspended in FACS buffer and stored on ice until staining.

### Flow cytometry

Cell suspensions were incubated with Zombie Aqua^TM^ dye (Fixable Viability kit, BioLegend 423102; 1:1000) for 20 minutes at room temperature in 1X PBS then washed with FACS Buffer (1X PBS, 2% FBS). Samples were incubated with Mouse BD Fc Block^TM^ (anti-CD16/CD32; BD Biosciences 553142; 1:10) for 10 minutes on ice then washed with FACS Buffer. Cell suspensions were stained for 20 minutes on ice in FACS buffer using tissue-specific antibody panels (**Supplemental Table 2**). Antibodies and dilutions are provided in **Supplemental Table 3**. Flow cytometry was performed using a BD FACS Symphony A1 Cell Analyzer, and data was analyzed using FlowJo V10.10.0 software. The gating strategies used to define cell types are outlined in the relevant figures (**Figure 1B**, **Figure 2A**, **Figure 3A**, and **Supplemental Figures 1, 2, 3, 5,** and **6**).

**Figure 1.**
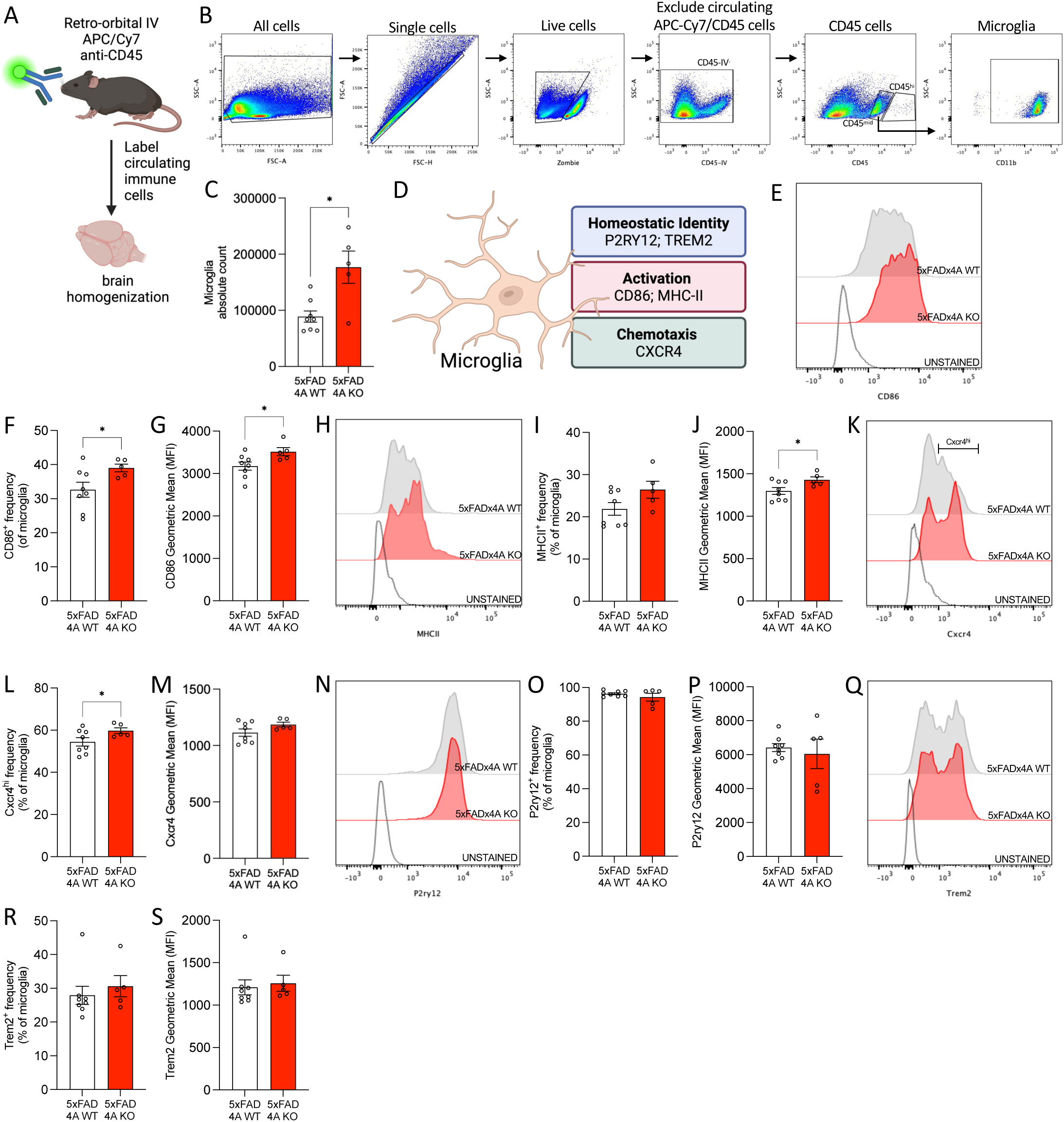
5xFAD Ms4a4a-deficient microglia assume a pro-inflammatory state. A. Schematic. Mice were retro-orbitally injected with an APC/Cy7-CD45 antibody to label circulating immune cells then brains were extracted and analyzed. B. Gating strategy to identify microglia in the brain. C. Microglia cell count. *, P = 0.034. D. Schematic of CD86, MHC class II, CXCR4, P2RY12, and TREM2 expression in microglia. E. Representative histogram of CD86 expression in microglia from 5xFAD 4A-WT (gray) and 5xFAD 4A-KO (red) mice. F. Quantification of CD86 frequency of expression in microglia. *, P = 0.027. G. Quantification of CD86 MFI in microglia. *, P = 0.032. H. Representative histogram of MHC class II expression in microglia from 5xFAD 4A-WT (gray) and 5xFAD 4A-KO (red) mice. I. Quantification of MHC class II frequency of expression in microglia. J. Quantification of MHC class II MFI in microglia. *, P = 0.028. K. Representative histogram of CXCR4 expression in microglia from 5xFAD 4A-WT (gray) and 5xFAD 4A-KO (red) mice. L. Quantification of CXCR4^hi^ frequency of expression in microglia. *, P = 0.05. M. Quantification of CXCR4 MFI in microglia. N. Representative histogram of P2RY12 expression in microglia from 5xFAD 4A-WT (gray) and 5xFAD 4A-KO (red) mice. O. Quantification of P2RY12 frequency of expression in microglia. P. Quantification of P2RY12 MFI in microglia. Q. Representative histogram of TREM2 expression in microglia from 5xFAD 4A-WT (gray) and 5xFAD 4A-KO (red) mice. R. Quantification of TREM2 frequency of expression in microglia. S. Quantification of TREM2 MFI in microglia. Data are representative of 5xFAD 4A-WT: n = 8; 5xFAD 4A-KO: n = 5; 6-month-old female mice. Each dot represents one animal. Graphs represent mean ± SEM. Statistical comparisons were made using an unpaired Student’s t-test with Welch’s correction. Outliers were identified and removed using the ROUT method (Q = 1%; R-S).

**Figure 2.**
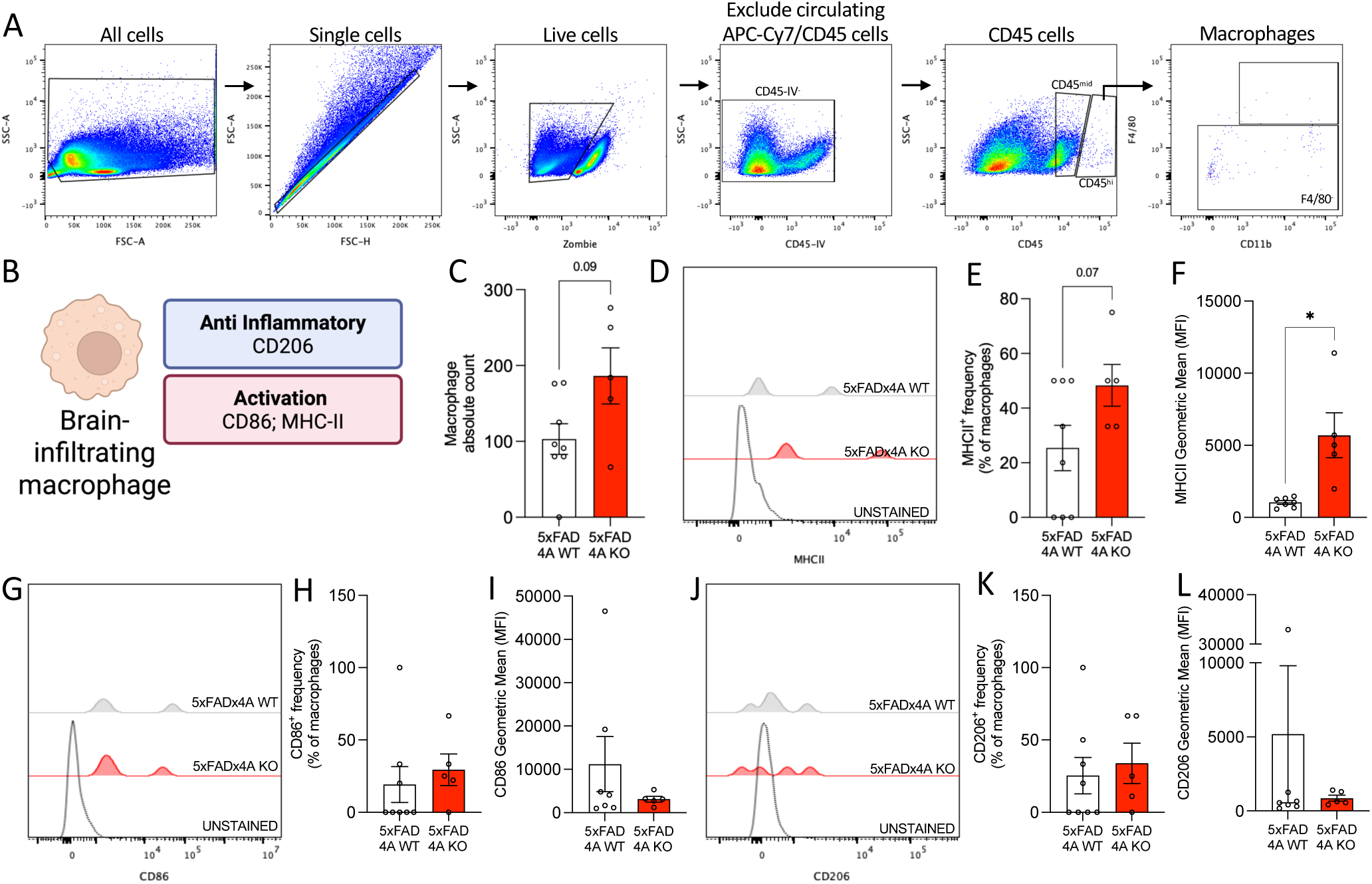
Ms4a4a loss selectively increases MHC class II expression in brain macrophages. A. Gating strategy to identify brain-infiltrating macrophages. B. Schematic of CD206, CD86, and MHC class II expression in brain-infiltrating macrophages. C. Brain-infiltrating macrophage cell count. D. Representative histogram of MHC class II expression in brain-infiltrating macrophages from 5xFAD 4A-WT (gray) and 5xFAD 4A-KO (red) mice. E. Quantification of MHC class II frequency of expression in brain-infiltrating macrophages. F. Quantification of MHC class II MFI in brain-infiltrating macrophages. *, P = 0.0398. G. Representative histogram of CD86 expression in brain-infiltrating macrophages from 5xFAD 4A-WT (gray) and 5xFAD 4A-KO (red) mice. H. Quantification of CD86 frequency of expression in brain-infiltrating macrophages. I. Quantification of CD86 MFI in brain-infiltrating macrophages. J. Representative histogram of CD206 expression in brain-infiltrating macrophages from 5xFAD 4A-WT (gray) and 5xFAD 4A-KO (red) mice. K. Quantification of CD206 frequency of expression in brain-infiltrating macrophages. L. Quantification of CD206 MFI in brain-infiltrating macrophages. Data are representative of 5xFAD 4A-WT: n = 8; 5xFAD 4A-KO: n = 5; 6-month-old female mice. Each dot represents one animal. Graphs represent mean ± SEM. Statistical comparisons were made using an unpaired Student’s t-test with Welch’s correction. Outliers were identified and removed using the ROUT method (Q = 1%; H-I, L).

**Figure 3.**
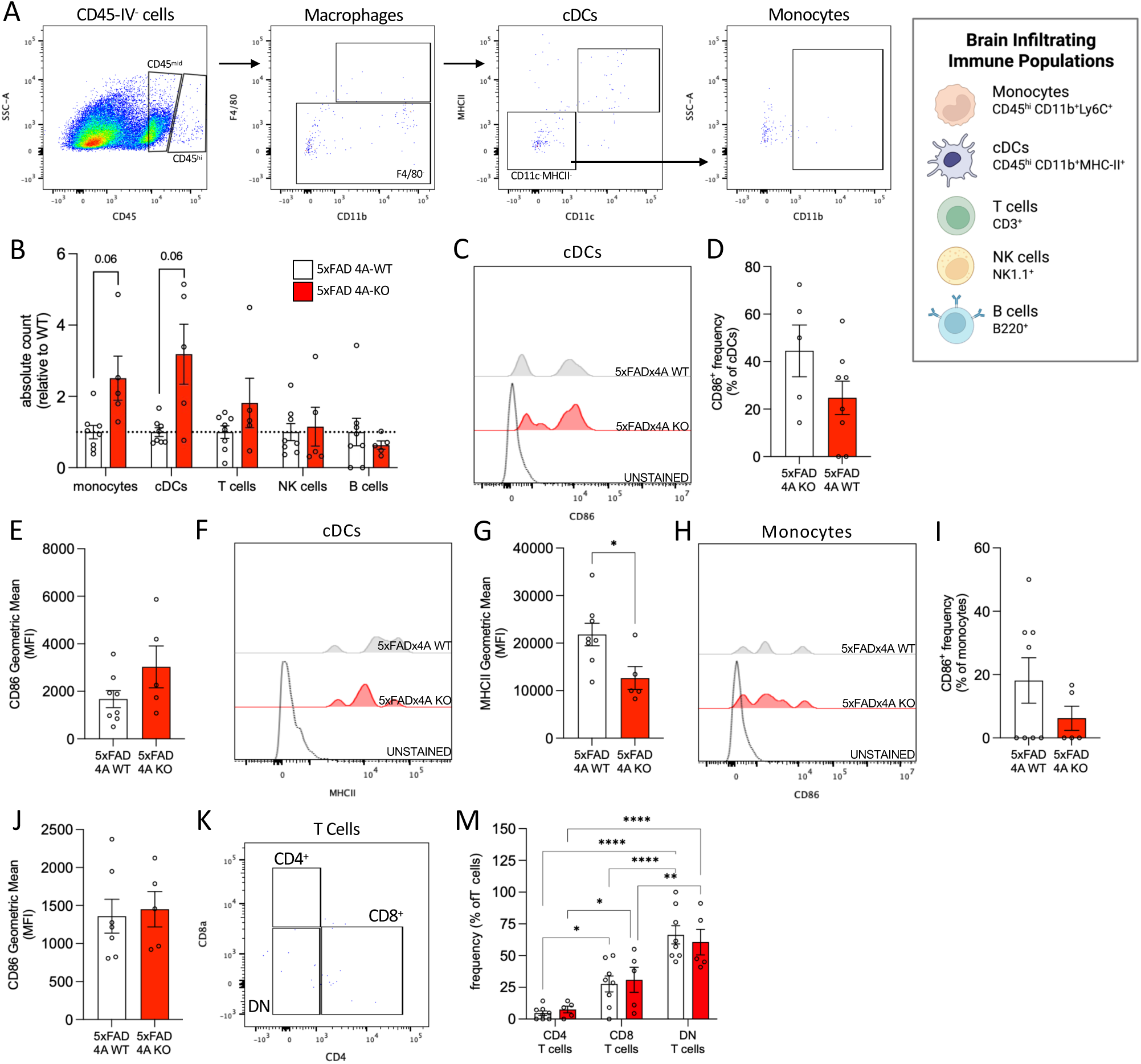
Brain-infiltrating monocytes and cDCs remain unaltered with loss of Ms4a4a in 5xFAD mice. A. Gating strategy to identify brain-infiltrating monocytes and cDCs. B. Brain-infiltrating immune cell counts in 5xFAD 4A-WT (white) and 5xFAD 4A-KO (red) mice, relative to WT. C. Representative histogram of CD86 expression in brain-infiltrating cDCs from 5xFAD 4A-WT (gray) and 5xFAD 4A-KO (red) mice. D. Quantification of CD86 frequency of expression in brain-infiltrating cDCs. E. Quantification of CD86 MFI in brain-infiltrating cDCs. F. Representative histogram of MHC class II expression in brain-infiltrating cDCs from 5xFAD 4A-WT (gray) and 5xFAD 4A-KO (red) mice. G. Quantification of MHC class II MFI in brain-infiltrating cDCs. P = 0.022. H. Representative histogram of CD86 expression in brain-infiltrating monocytes from 5xFAD 4A-WT (gray) and 5xFAD 4A-KO (red) mice. I. Quantification of CD86 frequency of expression in brain-infiltrating monocytes. J. Quantification of CD86 MFI in brain-infiltrating monocytes. K. Gating strategy to identify brain-infiltrating T-cell subtypes. M. Quantification of CD4, CD8, and DN T-cells in the brain. *, P < 0.05; **, P < 0.01; ****, P < 0.0001. Data are representative of 5xFAD 4A-WT: n = 8; 5xFAD 4A-KO: n = 5; 6-month-old female mice. Each dot represents one animal. Graphs represent mean ± SEM. Statistical comparisons were made using an unpaired Student’s t-test with Welch’s correction (B, D-E, G, I-J) and two-way ANOVA (M). Outliers were identified and removed using the ROUT method (Q = 1%; L).

### Statistical Analysis

Statistical analysis was performed in GraphPad Prism 11. Welch’s t-tests were used for pairwise comparisons between genotypes. Two-way ANOVA with multiple comparisons was applied to examine the effects of genotype, cell populations, and their interaction. *P <* 0.05 was considered statistically significant. Outliers were identified and removed using the ROUT method (Q = 1%).

## Results

### Ms4a4a loss increases microglial abundance and selectively remodels microglial activation in 5xFAD mice

We previously demonstrated that *Ms4a4a* loss reduces amyloid pathology and attenuates dense-core plaque-associated tau spread in 5xFAD mice [49, 50]. We further found that *Ms4a4a* deficiency alters activated microglial transcriptional states in the amyloid-bearing hippocampus [50]. To determine how *Ms4a4a* loss alters microglial phenotypes in the context of amyloid pathology, we performed flow cytometry on the brains of 6-month-old 5xFAD 4A-WT (n = 8) and 5xFAD 4A-KO (n = 5) mice (**Figure 1A**). Circulating immune cells were labeled *in vivo* by retro-orbital injection of an APC-Cy7/CD45 antibody and excluded during analysis. Brain-resident microglia were subsequently identified as expressing CD11b and an intermediate level of CD45 (termed: CD45^mid^)(**Figure 1B**). *Ms4a4a* loss significantly increased microglial counts in 5xFAD mice (**Figure 1C**).

Given our prior evidence that *Ms4a4a* loss alters activated microglial transcriptional states [50], we next asked whether increased microglial abundance was accompanied by changes in complementary features of microglial activation. We therefore examined markers associated with immune activation and costimulation (CD86), antigen presentation (MHC class II), and chemotactic signaling (CXCR4; **Figure 1D**). Loss of *Ms4a4a* increased both the frequency of CD86-positive microglia and CD86 mean fluorescence intensity (MFI; **Figure 1E–G**). MHC class II-positive microglial frequency did not differ significantly between genotypes, whereas MHC class II MFI was significantly increased in 5xFAD 4A-KO mice (**Figure 1H–J**). We also observed an increased frequency of CXCR4^hi^ microglia following *Ms4a4a* loss, while CXCR4 MFI remained unchanged (**Figure 1K–M**). Together, these findings indicate that *Ms4a4a* deficiency selectively increases both the prevalence and expression of activation-associated markers across distinct microglial populations.

We next assessed whether these activation-associated changes were accompanied by broader alterations in homeostatic or disease-associated microglial identity. The frequency and MFI of the homeostatic microglial marker P2RY12 was unchanged following *Ms4a4a* loss (**Figure 1N–P**). Trem2 is a regulator of the disease associated microglia state [55, 56]. Yet, neither the frequency of TREM2-positive microglia nor surface Trem2 MFI differed between 5xFAD 4A-WT and 5xFAD 4A-KO mice (**Figure 1Q–S**). Thus, *Ms4a4a* loss increases microglial abundance and selectively enhances CD86-, MHC class II-, and CXCR4-associated phenotypes without broadly altering P2RY12 or TREM2 expression.

### Ms4a4a loss selectively enhances MHC class II expression in brain macrophages

Peripheral myeloid cells can enter the brain during AD and contribute to inflammatory responses and disease pathology [13–20, 22]. Moreover, much of the functional characterization of *Ms4a4a* outside the CNS has focused on macrophages [42, 43, 45]. Considering the activation-associated changes observed in microglia, we asked whether *Ms4a4a* loss similarly alters the abundance or phenotype of brain-infiltrating macrophages in 5xFAD mice. Macrophages were identified by flow cytometry (**Figure 2A**), and markers of macrophage polarization and activation, including CD206, CD86, and MHC class II, were assessed (**Figure 2B**). Macrophage counts did not significantly differ between 5xFAD 4A-WT and 5xFAD 4A-KO mice (**Figure 2C**). We next examined macrophage marker expression. The frequency of MHC class II-positive macrophages was modestly higher in 5xFAD 4A-KO mice but did not reach statistical significance, whereas MHC class II MFI was significantly increased (**Figure 2D-F**). In contrast, neither the frequency nor MFI of CD86 differed between genotypes (**Figure 2G-I**). Expression of CD206 was similarly unchanged, with no differences in either the frequency of CD206-positive macrophages or CD206 MFI following *Ms4a4a* loss (**Figure 2J-L**). Together, these findings indicate that *Ms4a4a* deficiency selectively increases MHC class II expression in brain-infiltrating macrophages without broadly altering macrophage abundance or activation- or polarization-associated markers.

### Ms4a4a loss has limited effects on brain-infiltrating immune populations

Because macrophages represent only one peripheral immune population capable of entering the brain during AD, we next examined whether *Ms4a4a* loss altered other brain-infiltrating immune cells (**Figure 3A; Supplemental Figure 1**). Absolute counts of monocytes and conventional dendritic cells (cDCs) were modestly higher in 5xFAD 4A-KO mice but did not reach statistical significance (P = 0.06 for each), whereas T-cell, NK-cell, and B-cell counts were unchanged between genotypes (**Figure 3B**). Given the numerical increase in cDC abundance and their central role in antigen presentation, we tested whether *Ms4a4a* loss altered the activation phenotype of brain cDCs. CD86 frequency and MFI were unchanged between 5xFAD 4A-WT and 5xFAD 4A-KO mice (**Figure 3C-E**). In contrast, MHC class II MFI was significantly reduced in cDCs from 5xFAD 4A-KO mice (**Figure 3F-G**), indicating a selective effect of *Ms4a4a* loss on MHC class II expression rather than broader cDC activation. We next assessed activation-associated markers in brain-infiltrating monocytes. CD86-positive monocyte frequency and MFI were unchanged following *Ms4a4a* loss (**Figure 3H-J**). Thus, the modest numerical increase in brain monocytes was not accompanied by a detectable shift in the markers examined. Finally, we assessed the composition of brain-infiltrating T-cell populations (**Figure 3K**). The relative frequencies of CD4^+^, CD8^+^, and CD4/CD8 double-negative T-cells did not differ between 5xFAD 4A-WT and 5xFAD 4A-KO mice (**Figure 3M**). In both genotypes, CD8^+^ and double-negative T-cells represented a greater proportion of the T-cell compartment than CD4^+^ T-cells. Together, these findings demonstrate that *Ms4a4a* deficiency has limited effects on brain-infiltrating immune populations, with the most notable change being reduced MHC class II expression in cDCs.

### Ms4a4a loss produces limited, cell-type-selective changes in CNS-border immune compartments

Given the selective changes observed in brain macrophages and other infiltrating myeloid populations, we sought to determine whether *Ms4a4a* loss similarly altered immune populations within CNS-border compartments. We first examined the dural meninges, an immune-rich tissue positioned at the CNS interface that contributes to neuroimmune surveillance and inflammatory responses [15]. Flow cytometry was performed on dural meninges isolated from 5xFAD 4A-WT and 5xFAD 4A-KO mice (**Figure 4A, Supplemental Figure 2**). Absolute count of macrophages, monocytes, T-cells, and B-cells did not differ between genotypes, whereas cDC counts were modestly lower in 5xFAD 4A-KO mice but did not reach statistical significance (P = 0.06; **Figure 4B**). The relative frequencies of these populations within CD45-positive cells were similarly unchanged (**Figure 4C**).

**Figure 4.**
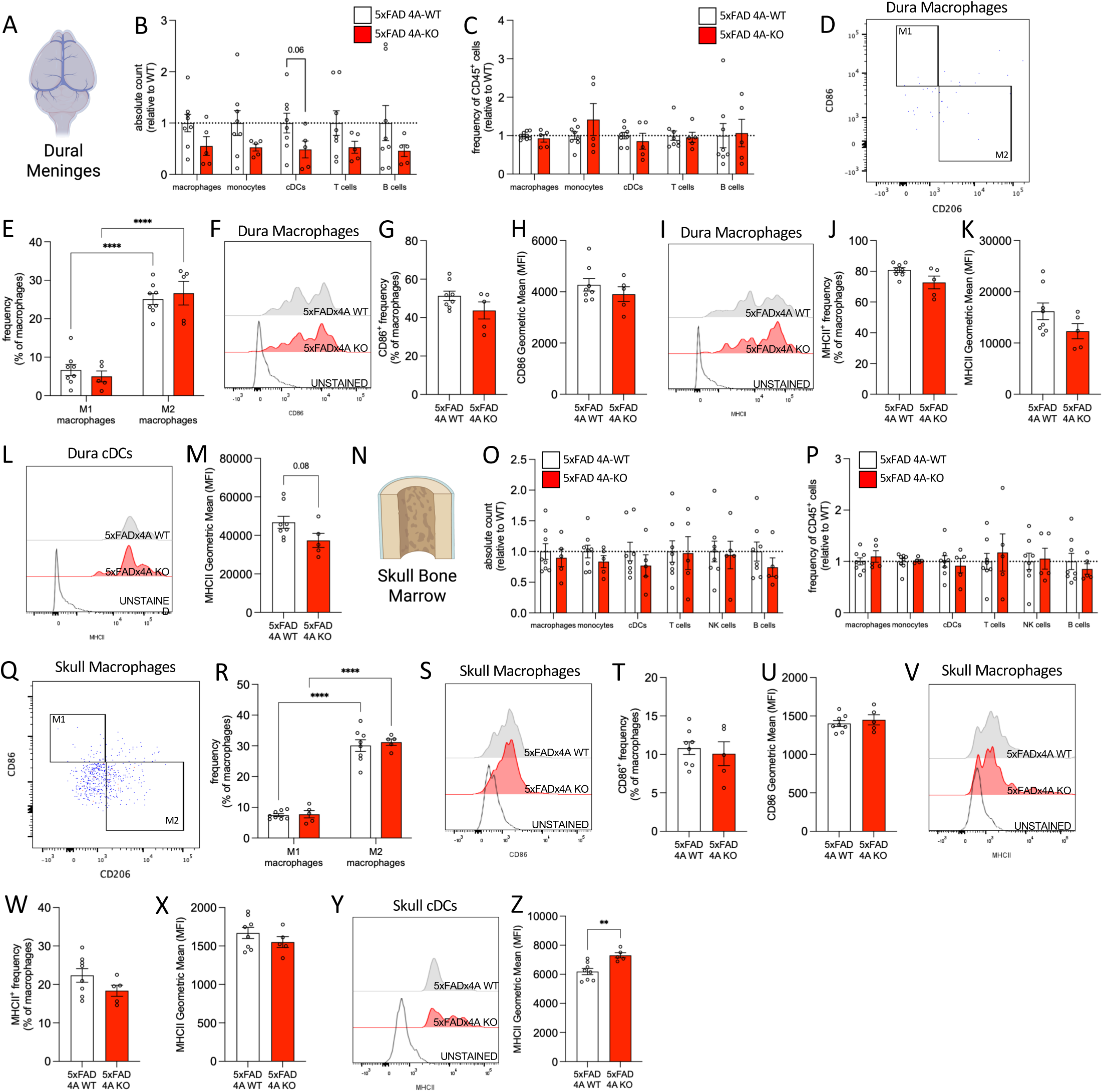
Macrophages in the dural meninges and skull bone marrow remain unaltered with the loss of Ms4a4a in 5xFAD mice. A. Schematic. Mice were retro-orbitally injected with an APC/Cy7-CD45 antibody to label circulating immune cells then dural meninges were extracted and analyzed. B. Dural meninges immune cell counts in 5xFAD 4A-WT (white) and 5xFAD 4A-KO (red) mice, relative to WT. C. Frequency of CD45^+^ cells by immune cell type in the dural meninges of 5xFAD 4A-WT (white) and 5xFAD 4A-KO (red) mice. D. Gating of dural macrophages to define M1 and M2 subpopulations. E. Frequency of M1-like and M2-like macrophages in the dural meninges of 5xFAD 4A-WT (white) and 5xFAD 4A-KO (red) mice. F. Representative histogram of CD86 expression in dural macrophages from 5xFAD 4A-WT (gray) and 5xFAD 4A-KO (red) mice. G. Quantification of CD86 frequency of expression in dural macrophages. H. Quantification of CD86 MFI in dural macrophages. I. Representative histogram of MHC class II expression in dural macrophages from 5xFAD 4A-WT (gray) and 5xFAD 4A-KO (red) mice. J. Quantification of MHC class II frequency of expression in dural macrophages. K. Quantification of MHC class II MFI in dural macrophages. L. Representative histogram of MHC class II expression in dural cDCs from 5xFAD 4A-WT (gray) and 5xFAD 4A-KO (red) mice. M. Quantification of MHC class II MFI in all dural cDCs. N. Schematic. Mice were retro-orbitally injected with an APC/Cy7-CD45 antibody to label circulating immune cells then skull bone marrow was extracted and analyzed. O. Skull bone marrow immune cell counts in 5xFAD 4A-WT (white) and 5xFAD 4A-KO (red) mice, relative to WT. P. Frequency of CD45^+^ cells by immune cell type in the skull bone marrow of 5xFAD 4A-WT (white) and 5xFAD 4A-KO (red) mice. Q. Gating of skull bone marrow macrophages to define M1 and M2 subpopulations. R. Frequency of M1-like and M2-like macrophages in the skull bone marrow of 5xFAD 4A-WT (white) and 5xFAD 4A-KO (red) mice. S. Representative histogram of CD86 expression in skull bone marrow macrophages from 5xFAD 4A-WT (gray) and 5xFAD 4A-KO (red) mice. T. Quantification of CD86 frequency of expression in skull bone marrow macrophages. U. Quantification of CD86 MFI in skull bone marrow macrophages. V. Representative histogram of MHC class II expression in skull bone marrow macrophages from 5xFAD 4A-WT (gray) and 5xFAD 4A-KO (red) mice. W. Quantification of MHC class II frequency of expression in skull bone marrow macrophages. X. Quantification of MHC class II MFI in skull bone marrow macrophages. Y. Representative histogram of MHC class II expression in skull bone marrow cDCs from 5xFAD 4A-WT (gray) and 5xFAD 4A-KO (red) mice. Z. Quantification of MHC class II MFI in all skull bone marrow cDCs. **, P = 0.002. Data are representative of 5xFAD 4A-WT: n = 8; 5xFAD 4A-KO: n = 5; 6-month-old female mice. Each dot represents one animal. Graphs represent mean ± SEM. Statistical comparisons were made using an unpaired Student’s t-test with Welch’s correction (B-C, G-H, J-K, N, P-Q, U-V, X-Y) and two-way ANOVA (E, S). ****, P < 0.0001.

We next examined whether *Ms4a4a* loss altered dural macrophage phenotypes (**Figure 4D**). CD206-positive, CD86-negative M2-like macrophages comprised a greater proportion of the macrophage population than CD86-positive, CD206-negative M1-like macrophages in both genotypes, with no genotype-dependent shift in their relative frequencies (**Figure 4E**). Consistent with this finding, neither the frequency nor MFI of CD86 or MHC class II differed between 5xFAD 4A-WT and 5xFAD 4A-KO mice (**Figure 4F–K**).

Because cDC abundance showed a modest genotype-associated difference, we next examined whether *Ms4a4a* loss altered the phenotype of dural cDCs. CD86 expression was unchanged, whereas MHC class II MFI was modestly lower in 5xFAD 4A-KO mice but did not reach statistical significance (P = 0.08; **Figure 4L-M; Supplemental Figure 2**). Among cDC subsets, cDC2 exhibited significantly higher MHC class II expression than cDC1 in 5xFAD 4A-KO mice, with a similar directional difference observed in 5xFAD 4A-WT mice (**Supplemental Figure 2**). Additional phenotyping of dural monocytes and T-cell subsets revealed no genotype-dependent differences (**Supplemental Figure 2**). Together, these findings indicate that *Ms4a4a* loss has limited effects on the overall composition and activation-associated phenotype of the dural immune compartment.

We next analyzed the skull bone marrow, a CNS-associated hematopoietic compartment that can supply immune cells to the meninges and brain [51, 53, 57]. Absolute counts of macrophages, monocytes, cDCs, T-cells, NK cells, and B-cells were unchanged following *Ms4a4a* loss, as were their relative frequencies within the CD45-positive cells (**Figure 4N-P; Supplemental Figure 3**). Similar to the dura, M2-like macrophages were more abundant than M1-like macrophages in both genotypes, with no genotype-dependent difference in their relative distribution (**Figure 4Q-R**). Consistent with this finding, neither CD86 nor MHC class II expression differed in skull bone marrow macrophages (**Figure 4S-X**).

Because cDCs exhibited selective phenotypic changes in other compartments, we next examined cDC marker expression in the skull bone marrow. CD86 expression was unchanged (**Supplemental Figure 3**), whereas MHC class II MFI was significantly increased in cDCs from 5xFAD 4A-KO mice (**Figure 4Y-Z**). MHC class II expression was higher in cDC2s than cDC1s in both genotypes, indicating a cDC subtype-associated difference that was independent of *Ms4a4a* genotype (**Supplemental Figure 3**). Monocyte activation markers and T-cell subset frequencies were similarly unchanged between genotypes (**Supplemental Figure 3**). Thus, *Ms4a4a* loss does not broadly alter the skull bone marrow immune compartment but selectively increases MHC class II expression in cDCs. Together, with the dural findings, these results indicate that *Ms4a4a* deficiency produces limited and cell type-specific changes within CNS-border immune compartments that are substantially more restricted than the microglial remodeling observed in the 5xFAD brain.

### Ms4a4a loss produces limited, compartment-specific changes in peripheral immune populations

Because *Ms4a4a* loss produced its strongest and most coordinated immune effects in microglia, with more limited changes in CNS-border compartments, we next explored whether these effects extended to peripheral immune tissues. We first examined the tibial bone marrow, a major hematopoietic compartment and source of circulating immune cells [22, 58]. Flow cytometry revealed no genotype-dependent differences in the absolute counts or relative frequencies of macrophages, monocytes, cDCs, T-cells, NK cells, or B-cells in 5xFAD 4A-WT and 5xFAD 4A-KO mice (**Figure 5A-C; Supplemental Figure 4**). We next examined the tibial bone marrow macrophage phenotype (**Figure 5D**). M2-like macrophages comprised a greater proportion of the macrophage compartment than M1-like macrophages in both genotypes, and the proportion of M2-like macrophages was significantly increased in 5xFAD 4A-KO mice (**Figure 5E**). Despite this shift, neither the frequency nor MFI of CD86 or MHC class II differed between genotypes (Figure 5F-K). Monocytes and cDCs similarly showed no genotype-dependent differences in CD86 and MHC class II expression (**Supplemental Figure 5**). MHC class II expression was higher in cDC2 than cDC1 cells in both genotypes, consistent with a cDC subtype-associated difference rather than an effect of *Ms4a4a* loss (**Supplemental Figure 5**). T-cell subset frequencies were also comparable between genotypes (**Figure 5L**). Thus, *Ms4a4a* loss largely preserves tibial bone marrow immune composition and myeloid marker expression despite a selective increase in M2-like macrophages.

**Figure 5.**
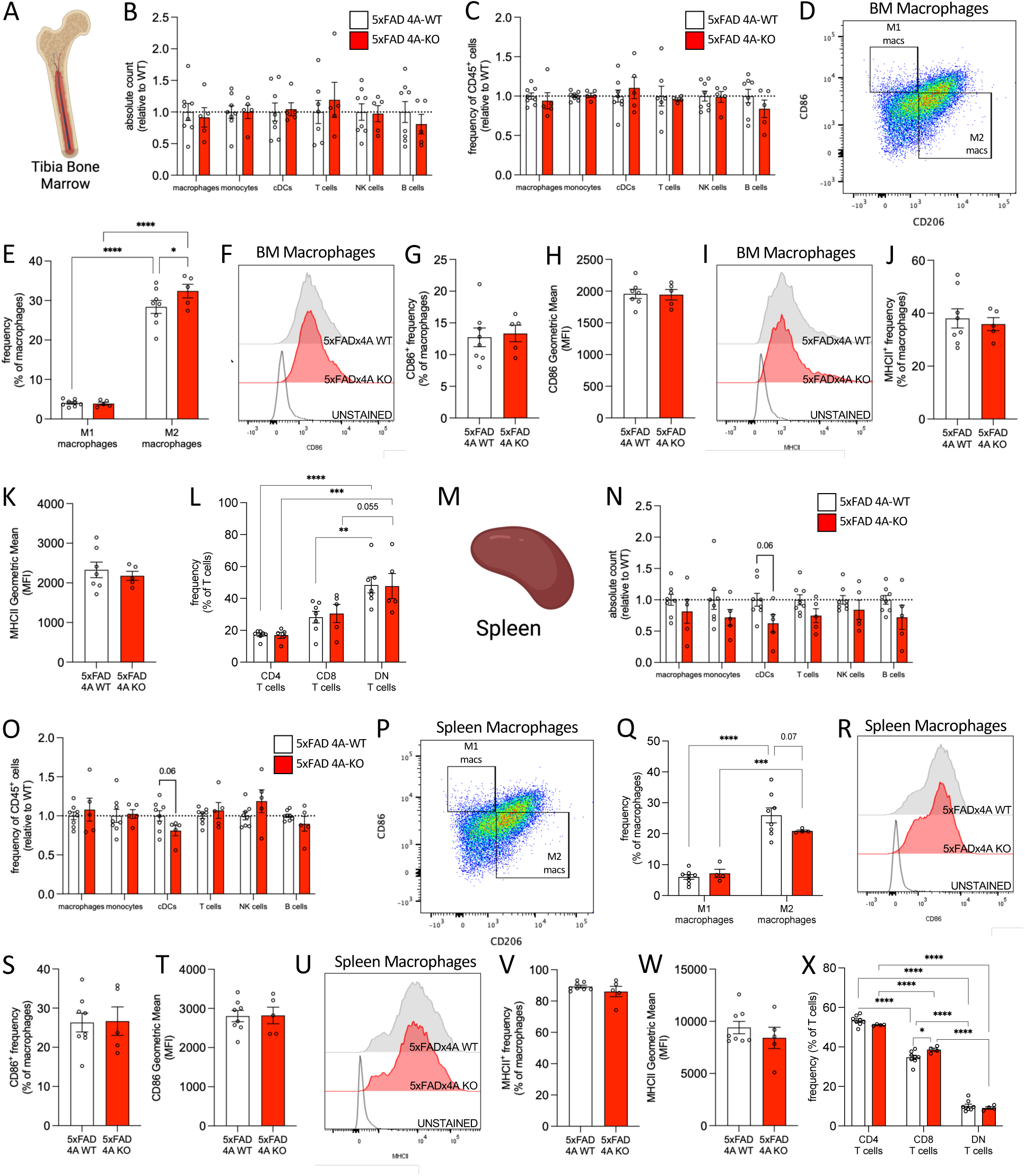
Loss of Ms4a4a in 5xFAD mice does not alter macrophages in the tibia bone marrow or spleen. A. Schematic. Mice were retro-orbitally injected with an APC/Cy7-CD45 antibody to label circulating immune cells then tibia bone marrow was extracted and analyzed. B. Tibia bone marrow immune cell counts in 5xFAD 4A-WT (white) and 5xFAD 4A-KO (red) mice, relative to WT. C. Frequency of CD45^+^ cells by immune cell type in the tibia bone marrow of 5xFAD 4A-WT (white) and 5xFAD 4A-KO (red) mice. D. Gating of tibia bone marrow macrophages to define subpopulations. E. Frequency of M1-like and M2-like macrophages in the tibia bone marrow of 5xFAD 4A-WT (white) and 5xFAD 4A-KO (red) mice. F. Representative histogram of CD86, CD206, and MHC class II expression in tibia bone marrow macrophages from 5xFAD 4A-WT (gray) and 5xFAD 4A-KO (red) mice. G. Quantification of CD86 frequency of expression in tibia bone marrow macrophages. H. Quantification of CD86 MFI in tibia bone marrow macrophages. I. Representative histogram of MHC class II expression in tibia bone marrow macrophages from 5xFAD 4A-WT (gray) and 5xFAD 4A-KO (red) mice. J. Quantification of MHC class II frequency of expression in tibia bone marrow macrophages. K. Quantification of MHC class II MFI in tibia bone marrow macrophages. L. Quantification of CD4, CD8, and DN T-cells in the tibia bone marrow. M. Schematic. Mice were retro-orbitally injected with an APC/Cy7-CD45 antibody to label circulating immune cells then spleens were extracted and analyzed. N. Splenic immune cell counts in 5xFAD 4A-WT (white) and 5xFAD 4A-KO (red) mice, relative to WT. O. Frequency of CD45^+^ cells by immune cell type in the spleens of 5xFAD 4A-WT (white) and 5xFAD 4A-KO (red) mice. P. Gating of splenic macrophages to define subpopulations. Q. Frequency of M1-like and M2-like macrophages in the spleens of 5xFAD 4A-WT (white) and 5xFAD 4A-KO (red) mice. R. Representative histogram of CD86 expression in splenic macrophages from 5xFAD 4A-WT (gray) and 5xFAD 4A-KO (red) mice. S. Quantification of CD86 frequency of expression in splenic macrophages. T. Quantification of CD86 MFI in splenic macrophages. U. Representative histogram of MHC class II expression in splenic macrophages from 5xFAD 4A-WT (gray) and 5xFAD 4A-KO (red) mice. V. Quantification of MHC class II frequency of expression in splenic macrophages. W. Quantification of MHC class II MFI in splenic macrophages. X. Quantification of CD4, CD8, and DN T-cells in the spleen. Data are representative of 5xFAD 4A-WT: n = 8; 5xFAD 4A-KO: n = 5; 6-month-old female mice. Each dot represents one animal. Graphs represent mean ± SEM. Statistical comparisons were made using an unpaired Student’s t-test with Welch’s correction (B-C, G-H, J-K, N-O, S-T, V-W) and two-way ANOVA (E, L, Q, X). *, P < 0.05; **, P < 0.01; ***, P < 0.001; ****, P < 0.0001. Outliers were identified and removed using the ROUT method (Q = 1%; B-C, L, Q, X).

We next assessed the spleen as a second peripheral immune compartment (**Figure 5M; Supplemental Figure 6**). Absolute counts and relative frequencies of macrophages, monocytes, T-cells, NK cells, and B-cells were unchanged following *Ms4a4a* loss. cDC abundance and frequency were modestly lower in 5xFAD 4A-KO mice but did not reach statistical significance (P = 0.06; **Figure 5N-O**). M2-like macrophages predominated over M1-like macrophages in both genotypes, with a modest reduction in the M2-like population in 5xFAD 4A-KO mice that also did not reach statistical significance (P = 0.07; **Figure 5P-Q**). CD86 and MHC class II expression were unchanged in splenic macrophages, monocytes, and cDCs (**Figure 5R-W; Supplemental Figure 7**). In contrast, analysis of splenic T-cell composition revealed a significant increase in the proportion of CD8+ T-cells in 5xFAD 4A-KO mice, whereas the remaining T-cell subsets were unchanged (**Figure 5X**). Together, these findings indicate that constitutive *Ms4a4a* loss does not broadly alter peripheral immune compartments in 5xFAD mice but produces selective, tissue-specific changes in macrophage and T-cell populations.

## Discussion

Our findings identify MS4A4A as a context-dependent regulator of immune cell state in the setting of amyloid pathology, with its most pronounced effects in microglia. Genetic studies first implicated the *MS4A* locus in AD risk and subsequently linked protective variation near *MS4A4A* to increased cerebrospinal fluid sTREM2, delayed age at onset, and altered microglial transcriptional states [6, 9, 36, 37, 39, 59]. More recent functional studies have implicated MS4A4A in the regulation of TREM2 signaling, microglial survival, phagocytosis, lysosomal function, and disease-associated microglia states [47]. Despite expression of *Ms4a4a* across multiple myeloid populations, we found that its loss produced a disproportionately strong effect on microglia, whereas other central, CNS-border, and peripheral immune populations showed more limited and heterogeneous responses. These findings extend prior work by demonstrating that the consequences of *Ms4a4a* loss are strongly dependent on cellular identity and tissue context.

Microglia adopt heterogeneous transcriptional and functional states in response to amyloid pathology, including disease-associated, interferon-responsive, lipid-associated, and antigen-presentation-associated states [55, 56, 60–63]. In our study, *Ms4a4a* loss increased microglial abundance and selectively altered CD86, MHC class II, and CXCR4 without changing P2RY12 or surface TREM2. Increased CD86 frequency and expression are consistent with enhanced immune activation and costimulatory capacity, while increased MHC class II expression suggests greater engagement of antigen-presentation-associated pathways. MHC class II-high microglial populations have been identified in mouse models of amyloidosis and human AD and appear to represent a distinct component of the microglial response rather than simply a loss of homeostatic identity [64–67]. For example, Aβ-specific CD4 T-cells can induce P2RY12-positive, MHC class II-positive microglia with enhanced Aβ uptake and plaque clearance in 5xFAD mice, whereas more recent studies have identified MHC class II-high microglia as part of an interferon-responsive state driven by interactions with infiltrating T-cells [68, 69]. These contrasting observations emphasize that MHC class II expression defines a microglial state but does not by itself establish whether that state is protective or detrimental. Together, preservation of P2RY12 and TREM2 alongside increased CD86 and MHC class II suggests that *Ms4a4a* loss selectively remodels activation- and antigen-presentation-associated features rather than driving a uniform transition from homeostatic to disease-associated microglia.

The CXCR4 phenotype further supports the selective expansion of a distinct microglial response. CXCR4 and its ligand CXCL12 regulate immune-cell migration and positioning [70–73], and CXCR4-dependent chemotaxis has been implicated in microglial responses to amyloid [55, 74, 75]. In human iPSC-derived microglia, loss of TREM2 impairs CXCL12/CXCR4-mediated migration and reduces microglial responses to Aβ pathology, linking this pathway to disease-relevant microglial positioning [76, 77]. Here, *Ms4a4a* deficiency increased the frequency of CXCR4-high microglia without altering CXCR4 MFI, suggesting enrichment of a CXCR4-expressing population rather than uniform induction of receptor expression. Notably, this occurred without changes in surface TREM2. Genetic and molecular studies have strongly linked MS4A4A to TREM2 biology, although the direction and nature of this relationship appear to depend on cellular context and the TREM2 species measured [36, 47]. Our findings therefore suggest that *Ms4a4a* loss can remodel microglial state and CXCR4-associated phenotypes *in vivo* without requiring a detectable change in steady-state surface TREM2 abundance. Whether TREM2 signaling, trafficking, or shedding is altered despite preserved surface expression remains to be determined.

These microglial changes provide additional cellular context for our previous observation that *Ms4a4a* deficiency enhances Aβ clearance, reduces plaque burden, increases plaque compaction, and limits dense-core plaque-associated tau spread [49, 50]. Importantly, these effects occur without increased microglial recruitment to plaques, distinguishing *Ms4a4a* from AD risk genes such as *TREM2*, for which impaired signaling substantially alters microglial survival, plaque association, and disease-associated state transitions [76, 78–84]. Our prior work also identified increased MMP-9 following *Ms4a4a* loss, providing one potential mechanism linking microglial remodeling to enhanced extracellular Aβ degradation. The current findings suggest that the coordinated microglial state induced by *Ms4a4a* deficiency may provide the cellular setting in which these protective pathological effects occur. However, CD86, MHC class II, and CXCR4 expression do not establish the underlying functional mechanism, and additional studies will be required to determine which components of this microglial response are necessary for enhanced Aβ clearance and reduced downstream tau pathology.

Other myeloid populations within the brain exhibited considerably more restricted responses to *Ms4a4a* loss. Brain macrophage abundance and CD206 and CD86 expression were unchanged, whereas MHC class II expression was increased. Macrophages recruited to or residing within the diseased CNS can acquire transcriptional and antigen-presentation programs that differ substantially from those of circulating or tissue-resident peripheral macrophages [85, 86]. Our findings similarly suggest that *Ms4a4a* loss does not induce a broad macrophage activation program but selectively modifies an antigen-presentation-associated feature within the brain environment. Brain cDCs showed the opposite pattern, with reduced MHC class II expression despite a modest numerical increase. Dendritic cells provide an important interface between innate and adaptive immunity, and MHC class II and CD86 expression are central to their capacity to activate antigen-specific T-cells [87–89]. In AD models, dendritic cells have been identified in perivascular and meningeal regions containing Aβ and can participate in the regulation of Aβ-specific T-cell responses [90–93]. Thus, MHC class II emerges as a recurrent but cell type-dependent feature of *Ms4a4a* loss: increased in microglia and brain macrophages but reduced in brain cDCs. These divergent responses argue against a uniform effect on antigen presentation and instead suggest that MS4A4A regulates MHC class II-associated phenotypes in a cell- and context-dependent manner.

The relative preservation of the dura and skull bone marrow provides further evidence for this compartment specificity. The meninges and adjacent skull marrow are increasingly recognized as integrated components of CNS immune surveillance, with skull marrow supplying immune cells directly to the meninges through specialized vascular channels and responding to neurological disease, including AD [51, 53, 93–98]. Despite this close anatomical relationship to the brain, overall immune composition in the dura and skull marrow was largely preserved following *Ms4a4a* loss. This suggests that the prominent microglial phenotype does not arise as part of parallel, large-scale remodeling of adjacent CNS-border immune reservoirs. Nevertheless, selective effects were evident in cDCs, including increased MHC class II expression in skull marrow cDCs and a more modest reduction in dural cDC MHC class II. These findings further reinforce our conclusion that *Ms4a4a* loss does not produce a uniform inflammatory response across anatomically connected compartments but instead modifies selected immune phenotypes in a tissue- and cell type-dependent manner.

Peripheral immune compartments were similarly resistant to broad remodeling following *Ms4a4a* loss. Immune-cell abundance and most activation-associated markers were unchanged in the tibial bone marrow and spleen, although M2-like macrophages were increased in tibial marrow and splenic CD8 T-cell frequency was increased. MS4A4A has been characterized in peripheral macrophages, where its expression is associated with monocyte-to-macrophage differentiation, tissue-resident or alternatively activated macrophage states, and Dectin-1-dependent signaling [43, 45]. The absence of a common macrophage phenotype across dura, skull marrow, tibial marrow, spleen, and brain therefore underscores the importance of tissue context in determining the consequences of *Ms4a4a* loss. The splenic CD8 T-cell finding may also be relevant given increasing evidence that adaptive immune responses contribute to amyloid pathology and that CD8 T-cells can interact with microglia in the AD brain [15, 19, 69, 99–102]. However, whether the modest peripheral T-cell change influences CNS pathology or represents an independent systemic consequence of *Ms4a4a* deficiency remains unknown. Overall, the peripheral findings argue against a coordinated systemic immune phenotype despite established roles for MS4A4A in peripheral myeloid cells.

Together, our findings identify MS4A4A as a cell type- and tissue-dependent regulator of immune responses to amyloid pathology. Although *Ms4a4a* is expressed across multiple myeloid populations, its loss produces a coordinated microglial response with comparatively limited and heterogeneous effects across other central and peripheral immune compartments. Combined with genetic evidence linking the *MS4A* locus to AD risk and resilience and emerging studies implicating MS4A4A in TREM2 signaling, phagocytosis, and lysosomal function, these findings support MS4A4A-dependent pathways as potential targets for modulating disease-relevant microglial responses. Defining which components of the *Ms4a4a*-deficient microglial state contribute to enhanced Aβ clearance and reduced downstream tau pathology will be important for determining whether these protective pathways can be selectively harnessed without broadly disrupting systemic immunity.

## List of Abbreviations

AD: Alzheimer’ Disease
Aβ: amyloid beta
cDC: conventional dendritic cell
CNS: Central Nervous System
MFI: mean fluorescence intensity
sTREM2: soluble TREM2

## Acknowledgments

We thank Alector for providing the *Ms4a4a* KO mice used in this study. We thank Laura Piccio, Samantha Swift, Jasmin Herz, and Torri Ball for thoughtful discussions. This work was supported by access to equipment made possible by the Hope Center for Neurological Disorders, the Neurogenomics and Informatics Center, and the Departments of Neurology and Psychiatry at Washington University School of Medicine. Diagrams were generated using BioRender.com. This manuscript is the result of funding in whole or in part by the National Institutes of Health (NIH). It is subject to the NIH Public Access Policy. Through acceptance of this federal funding, NIH has been given a right to make this manuscript publicly available in PubMed Central upon the Official Date of Publication, as defined by NIH. The content is solely the responsibility of the authors and does not necessarily represent the official views of the National Institutes of Health.

## Funding

Funding provided by the National Institutes of Health (AG062734), Hope Center for Neurological Disorders, Chan Zuckerberg Initiative (CMK), Thome Memorial Foundation, Alzheimer’s Association, and BrightFocus Foundation.

## Authors’ Contributions

Designed experiments: EPD, CMK. Performed and analyzed experiments: EPD, ACV, AKI, CMK. Provided funding: CMK. Wrote the manuscript: EPD, CMK. Revised and approved manuscript: EPD, ACV, AKI, CMK.

## Ethics approval and consent to participate

Animal care and surgical procedures were approved by the Animal Studies Committee of Washington University School of Medicine (Protocol #23-0237) in accordance with guidelines from the United States National Institutes of Health. This work did not involve the use of material from human subjects.

## Consent for publication

All authors have read and approved the final manuscript for publication.

## Availability of data and materials

All data generated or analyzed during this study are included in this published article and its supplementary information files.

## Competing interests

The authors declare that they have no competing interests.

## Notes

### Competing Interest Statement

The authors have declared no competing interest.

