## Supplemental Figures for "Ms4a4a loss preferentially remodels microglia with limited, compartment-specific effects across central and peripheral immune compartments in 5xFAD mice"

**Supplemental Figure Legends**

**Supplemental Figure 1. Gating strategy for identifying lymphoid cells in the brain.** Gating strategy used to identify B-cells and CD4, CD8, and DN T-cells in the brain.

**Supplemental Figure 2. Loss of *Ms4a4a* has minimal effect on immune landscape of the dura of 5xFAD mice.** A. Gating strategy used to identify macrophages, monocytes, and cDCs in the dural meninges. B. Gating strategy used to identify B-cells and CD4, CD8, and DN T-cells in the dural meninges. C-J. Characterization of immune cells in the dural meninges. C. Representative histogram of CD86 expression in dural monocytes from 5xFAD 4A-WT (gray) and 5xFAD 4A-KO (red) mice. D. Quantification of CD86 frequency of expression in dural monocytes. E. Quantification of CD86 MFI in dural monocytes. F. Representative histogram of CD86 expression in dural cDCs from 5xFAD 4A-WT (gray) and 5xFAD 4A-KO (red) mice. G. Quantification of CD86 frequency of expression in dural cDCs. H. Quantification of CD86 MFI in dural cDCs. I. Quantification of MHC class II MFI in dural cDC1s and cDC2s. *, $P = 0.0497$. J. Quantification of CD4, CD8, and DN T-cells in dural meninges. Data are representative of 5xFAD 4A-WT: $n = 8$; 5xFAD 4A-KO: $n = 5$; 6-month-old female mice. Each dot represents one animal. Graphs represent mean ± SEM. Statistical comparisons were made using an unpaired Student's t-test with Welch's correction (D-E, G-H) and two-way ANOVA (I-J). *, $P < 0.05$; ***, $P < 0.001$; ****, $P < 0.0001$.

**Supplemental Figure 3. Loss of *Ms4a4a* has minimal effect on immune landscape of the skull bone marrow of 5xFAD mice.** A. Gating strategy used to identify macrophages, monocytes, and cDCs in the skull bone marrow. B. Gating strategy used to identify B-cells and CD4, CD8, and DN T-cells in the skull bone marrow. C. Representative histogram of CD86 expression in skull bone marrow monocytes from 5xFAD 4A-WT (gray) and 5xFAD 4A-KO (red) mice. D. Quantification of CD86 frequency of expression in skull bone marrow monocytes. E. Quantification of CD86 MFI in skull bone marrow monocytes. F. Representative histogram of CD86 expression in skull bone marrow cDCs from 5xFAD 4A-WT (gray) and 5xFAD 4A-KO (red) mice. G. Quantification of CD86 frequency of expression in skull bone marrow cDCs. H. Quantification of CD86 MFI in skull bone marrow cDCs. I. Quantification of MHC class II MFI in skull bone marrow cDC1s and cDC2s. J. Quantification of CD4, CD8, and DN T-cells in skull bone marrow. Data are representative of 5xFAD 4A-WT: n = 8; 5xFAD 4A-KO: n = 5; 6-month-old female mice. Each dot represents one animal. Graphs represent mean ± SEM. Statistical comparisons were made using an unpaired Student's t-test with Welch's correction (D-E, G-H) and two-way ANOVA (I-J). *, $P < 0.05$; **, $P < 0.01$; ****, $P < 0.0001$.

**Supplemental Figure 4. Gating strategy for identifying immune cells in the tibial bone marrow.** A. Gating strategy used to identify macrophages, monocytes, and cDCs in the tibial bone marrow. B. Gating strategy used to identify B-cells and CD4, CD8, and DN T-cells in the tibial bone marrow.

**Supplemental Figure 5. *Ms4a4a* deficiency has a minimal effect on the immune landscape of the tibial bone marrow of 5xFAD mice.** A. Schematic of tibia bone marrow. B. Representative histogram of CD86 expression in tibia bone marrow monocytes from 5xFAD 4A-WT (gray) and 5xFAD 4A-KO (red) mice. C. Quantification of CD86 frequency of expression in tibia bone marrow monocytes. D. Quantification of CD86 MFI in tibia bone marrow monocytes. E. Representative histogram of MHC class II expression in tibia bone marrow monocytes from 5xFAD 4A-WT (gray) and 5xFAD 4A-KO (red) mice. F. Quantification of MHC class II frequency of expression in tibia bone marrow monocytes. G. Quantification of MHC class II MFI in tibia bone marrow monocytes. H. Representative histogram of CD86 expression in tibia bone marrow cDCs from 5xFAD 4A-WT (gray) and 5xFAD 4A-KO (red) mice. I. Quantification of CD86 frequency of expression in tibia bone marrow cDCs. J. Quantification of CD86 MFI in tibia bone marrow cDCs. K. Representative histogram of MHC class II expression in tibia bone marrow cDCs from 5xFAD 4A-WT (gray) and 5xFAD 4A-KO (red) mice. L. Quantification of MHC class II MFI in all tibia bone marrow cDCs. M. Quantification of MHC class II MFI in tibia bone marrow cDC1s and cDC2s. ****, $P < 0.0001$. Data are representative of 5xFAD 4A-WT: n = 8; 5xFAD 4A-KO: n = 5; 6-month-old female mice. Each dot represents one animal. Graphs represent mean ± SEM. Statistical comparisons were made using an unpaired Student's t-test with Welch's correction (C-D, F-G, I-J, L) and two-way ANOVA (M).

**Supplemental Figure 6. Gating strategy for identifying immune cells in the spleen.** A. Gating strategy used to identify macrophages, monocytes, and cDCs in the spleen. B. Gating strategy used to identify B-cells and CD4, CD8, and DN T-cells in the spleen.

**Supplemental Figure 7. *Ms4a4a* deficiency has a minimal effect on the immune landscape of the spleen of 5xFAD mice.** A. Schematic of the spleen. B. Representative histogram of CD86 expression in splenic monocytes from 5xFAD 4A-WT (gray) and 5xFAD 4A-KO (red) mice. C. Quantification of CD86 frequency of expression in splenic monocytes. D. Quantification of CD86 MFI in splenic monocytes. E. Representative histogram of MHC class II expression in splenic monocytes from 5xFAD 4A-WT (gray) and 5xFAD 4A-KO (red) mice. F. Quantification of MHC class II frequency of expression in splenic monocytes. G. Quantification of MHC class II MFI in splenic monocytes. H. Representative histogram of CD86 expression in splenic cDCs from 5xFAD 4A-WT (gray) and 5xFAD 4A-KO (red) mice. I. Quantification of CD86 frequency of expression in splenic cDCs. J. Quantification of CD86 MFI in splenic cDCs. K. Representative histogram of MHC class II expression in splenic cDCs from 5xFAD 4A-WT (gray) and 5xFAD 4A-KO (red) mice. L. Quantification of MHC class II frequency of expression in all splenic cDCs. M. Quantification of MHC class II MFI in splenic cDC1s and cDC2s. Data are representative of 5xFAD 4A-WT: n = 8; 5xFAD 4A-KO: n = 5; 6-month-old female mice. Each dot represents one animal. Graphs represent mean ± SEM. Statistical comparisons were made using an unpaired Student's t-test with Welch's correction (C-D, F-G, I-J, L) and two-way ANOVA (M).

### Supplemental Figure 1

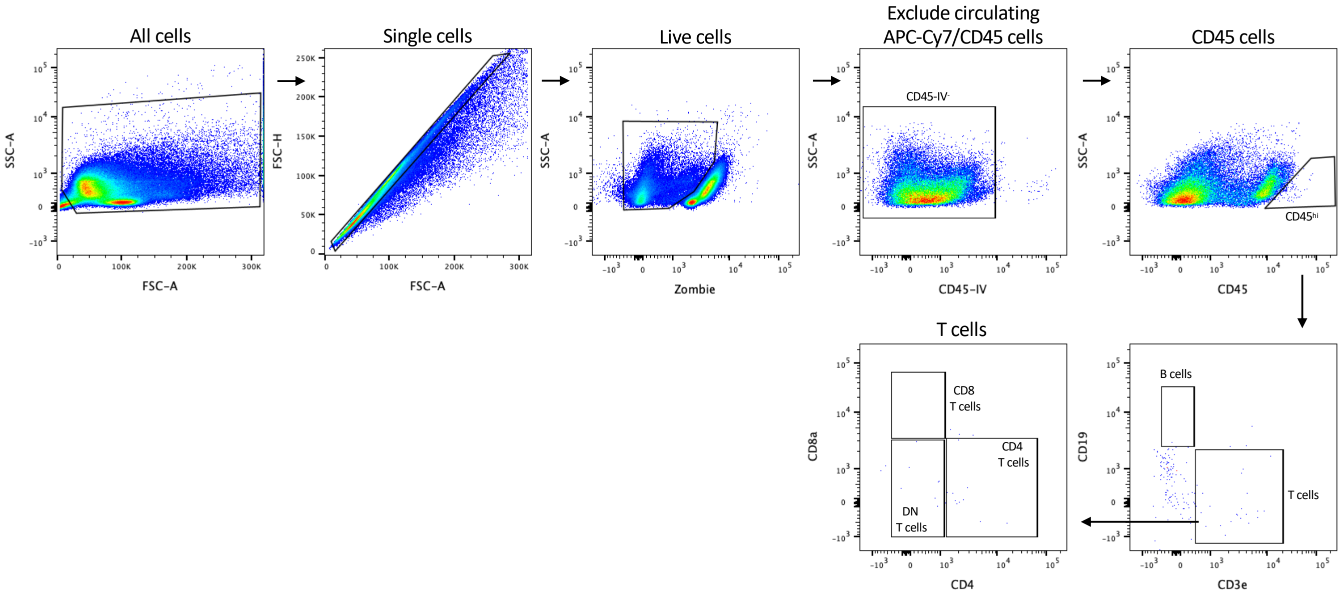

### Supplemental Figure 2

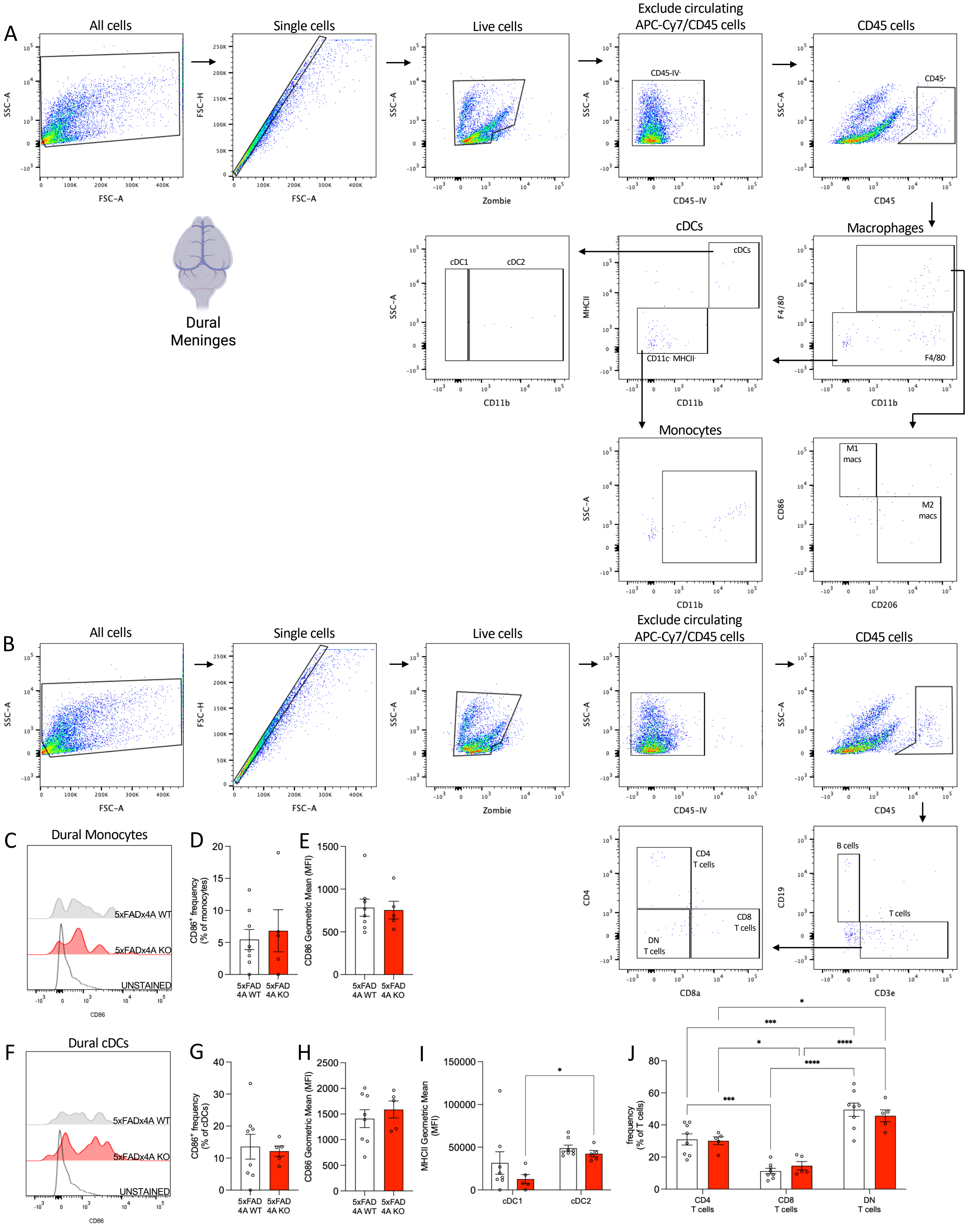

### Supplemental Figure 3

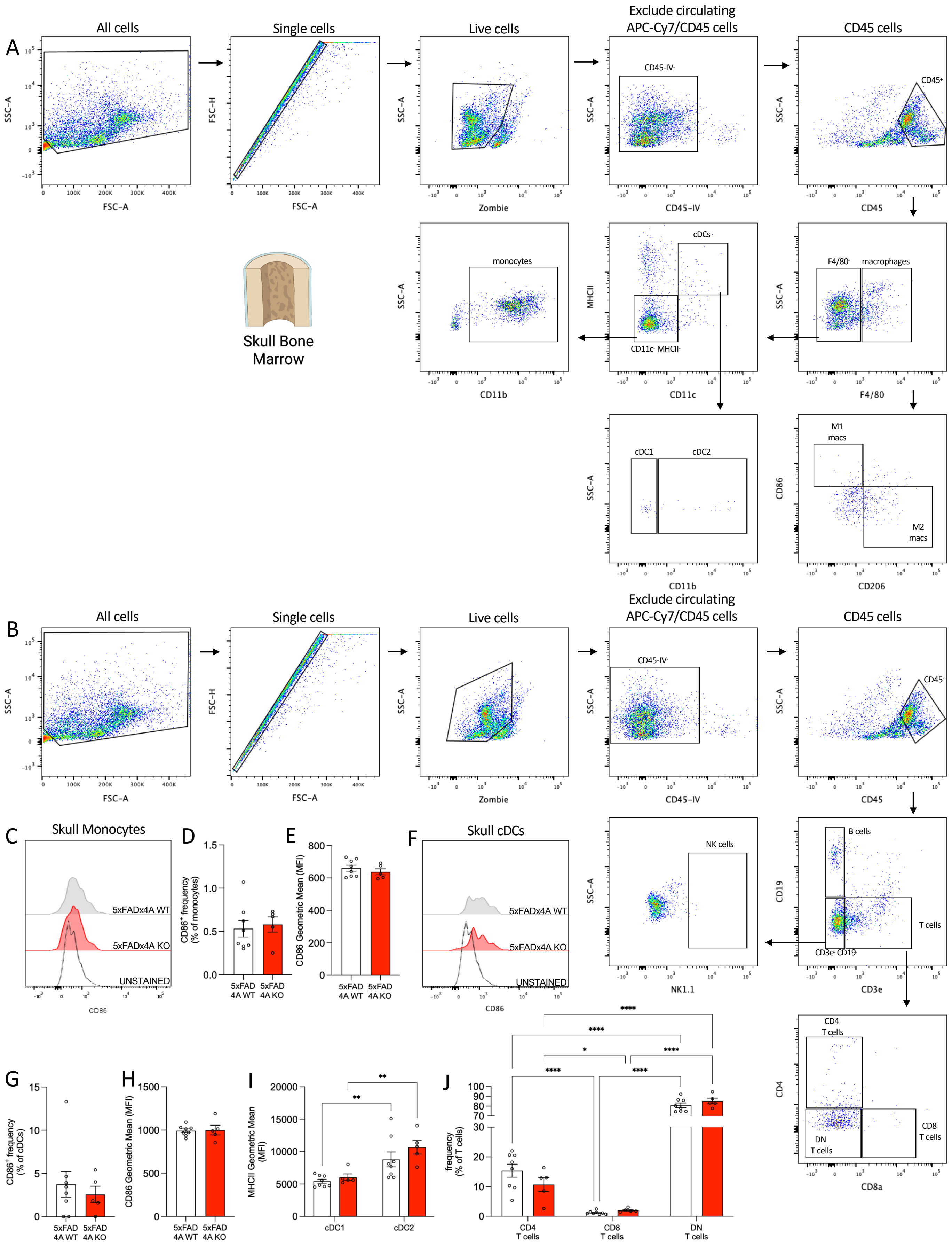

### Supplemental Figure 4

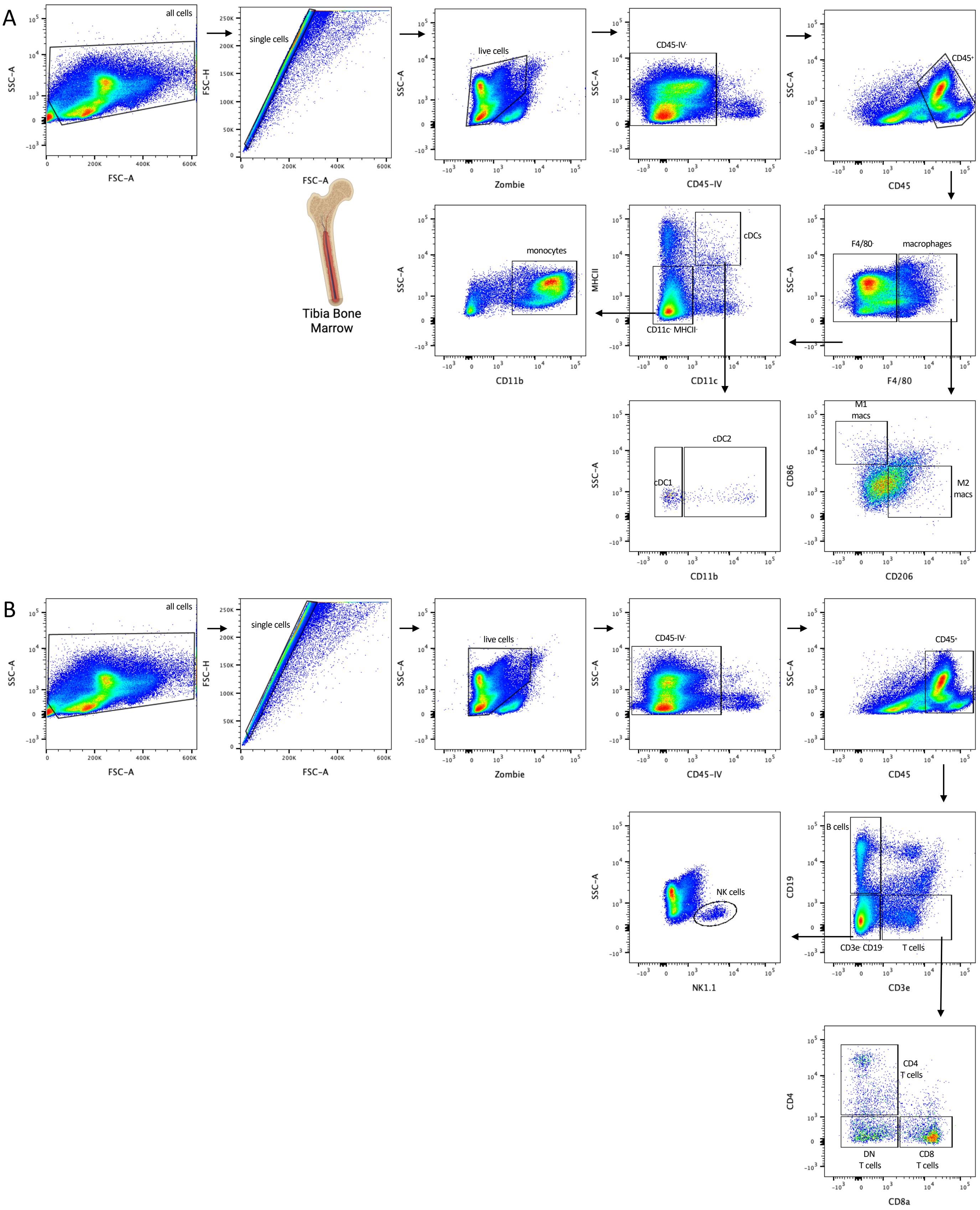

### Supplemental Figure 5

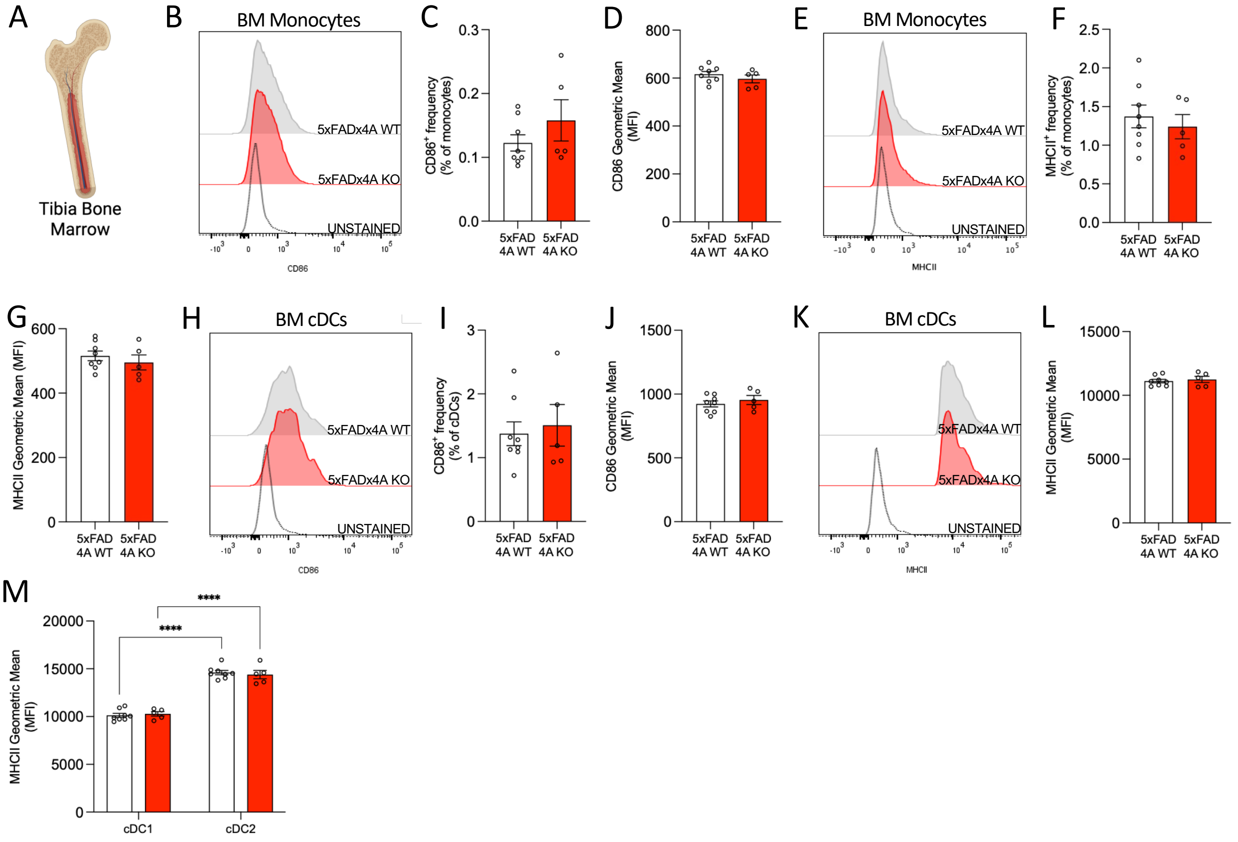

### Supplemental Figure 6

A

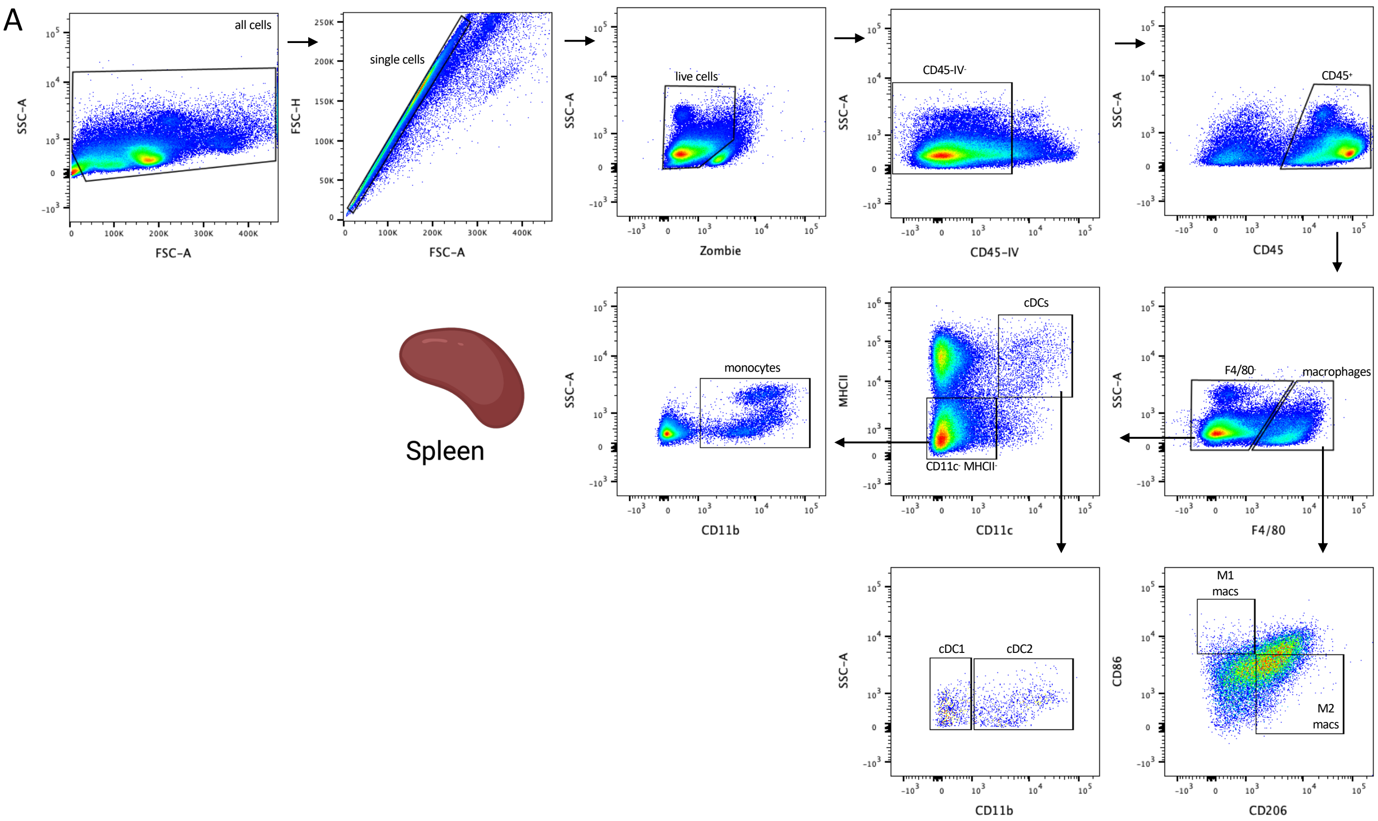


B

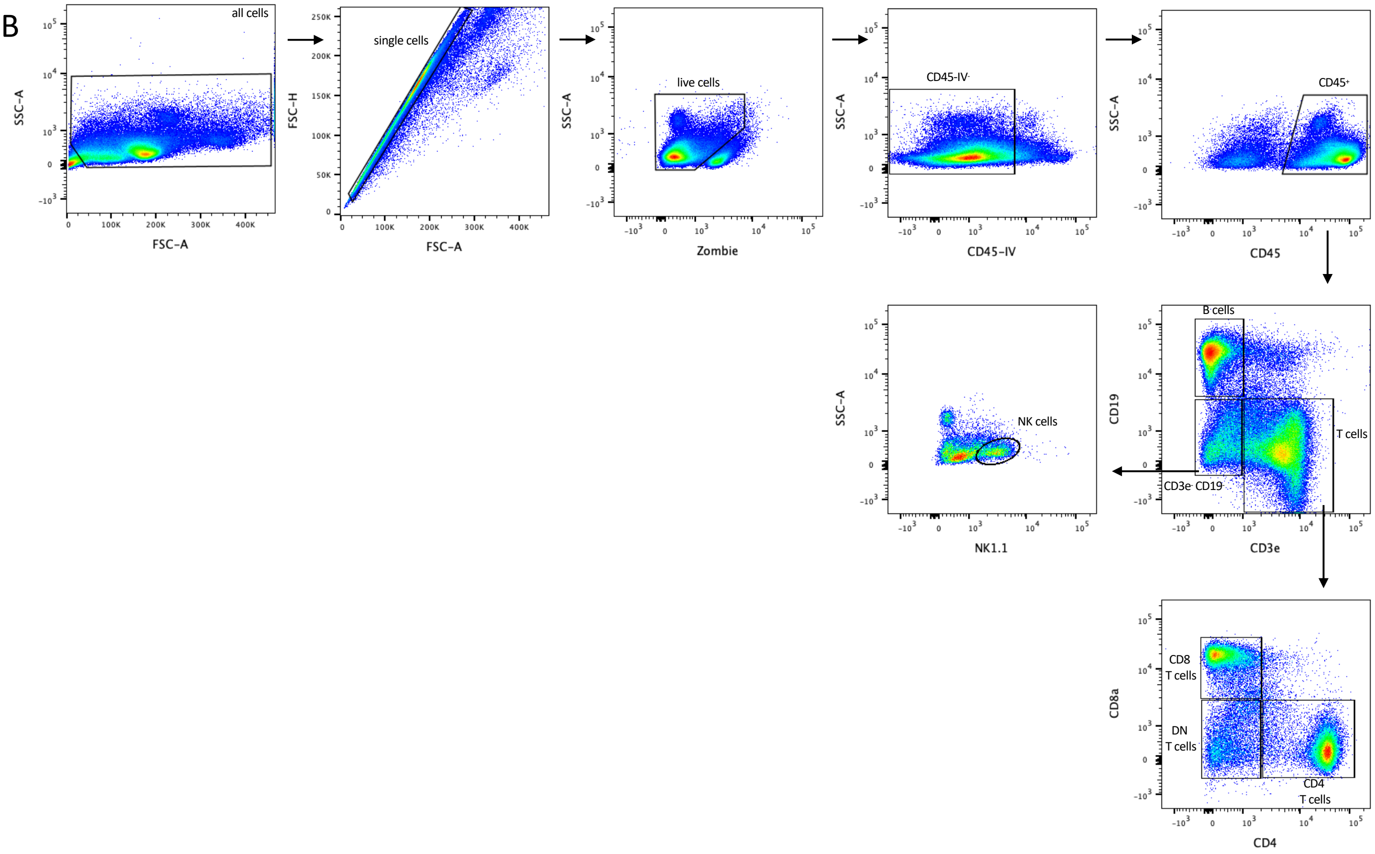

### Supplemental Figure 7

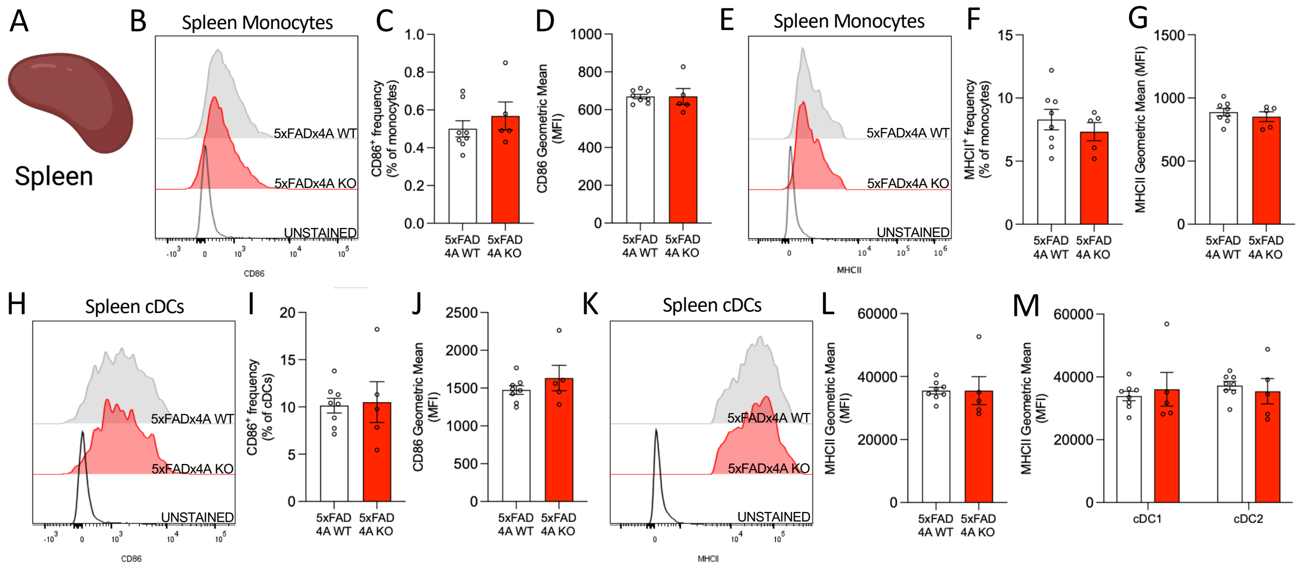
